# Interactions of Phototropism and Gravitropism in Cyanobacteria

**DOI:** 10.64898/2026.02.21.707229

**Authors:** Colin Gates, Haridas Mundoor, Ivan Smalyukh, Jeffrey C. Cameron

## Abstract

While gene expression in bacteria has been shown to be affected by near-zero or extremely high gravity, a mechanism has not been established to date. In larger organisms, gravity-sensing mechanisms usually rely on a dense body applying directional pressure which can be detected by the cell. Herein we demonstrate a means of observing the effect of gravity on cyanobacteria by differential expression of native pigments in response to both gravity and light. We observe that in the cyanobacterium *Synechococcus* sp. PCC 7002, the distribution of pigmentation within the cell, and across cell colonies, is regulated by combined directional sensing of incoming light, adhesion to a surface via extracellular matrix, and applied external force, including the normal force of gravity applied to the cell. Cells grown on a substrate orient their thylakoids on the cell faces proximal and distal to the substrate and locate both chlorophyll and phycobilins in both of these membrane regions; phycobilins are primarily targeted to the membrane region nearest to the light source, while chlorophyll is preferentially expressed in the region opposite the overall external force applied to the cell. The mechanism for distribution of pigments appears to be regulated by presence of polyphosphate bodies within the cell, and removal of polyphosphate negates the cell’s ability to sense external forces. Furthermore, colonial morphology is affected by application of force, with cells responding to the secretions of other cells along a gradient along the expected response to shading. These results represent a critical step toward understanding basal phototrophic regulatory mechanisms of light use and demonstrate the first known intracellular directional gravity response mechanism in a prokaryote.

**Statement of Significance:** To date, no directionally sensitive gravity response mechanism has been observed in any prokaryote. We demonstrate the first evidence of a directional response to external force in a cyanobacterium. This pigment distribution force-directed response interacts with the conventional response to directional light. Furthermore, the cells appear to be able to respond to the presence of other cells above them via intercellular signaling which is not simply due to shading by light.

## Introduction

Gene expression in bacteria has previously been shown to be affected by near-zero or extremely high gravity (1,2). Cyanobacteria in particular have been recognized as suffering impaired growth as a result of the loss of consistent, directional, Earth-like gravity (3). A wide range of effects on motility have been noted since the development of artificial microgravity systems (4), but these impacts are ascribed to a loss of gravity in general rather than a loss of directional sensitivity to gravity in bacteria. In contrast, animal and plant cells are known to have specific consequences of the loss of gravitational force applied in a specific direction (5-7). In larger organisms, gravity-sensing mechanisms typically consist of a dense body applying directional pressure to a sensitive region of the cell, such as the plant amyloplast system (8), strain on the mammalian cytoskeleton from attached organelles (9), or fungal protein crystals (10). However, no mechanistic basis for directional gravitropic sensing behavior has been established in bacteria previously, to the best of our knowledge.

Cyanobacteria are among the fastest-growing photosynthetic organisms, capable of achieving doubling times as short as 2 hours under ideal conditions (11). They are extremely resilient, capable of colonizing almost every environment on the planet, and play crucial roles in global nutrient cycling, especially in their capacity as nitrogen fixers (12). Additionally, cyanobacteria share a common ancestor with the chloroplasts of all photosynthetic eukaryotes, including plants, which were acquired via the endosymbiosis of cyanobacteria by an early eukaryote hundreds of millions of years ago (13). Thus, they are common model organisms for understanding plant photosynthesis, which developed from the cyanobacterial process. As natural producers of a range of high-value biological compounds such as pigments and lipids, cyanobacteria are commonly used in the bioproduction and animal feed industries (14). Their ability to tolerate harsh conditions, together with their growth rate, makes cyanobacteria an ideal source of biomass and oxygen in spaceflight and in extraplanetary endeavors (15,16).

While some cyanobacteria are naturally planktonic and preferentially inhabit liquid environments, many strains will aggregate into colonies and form biofilms if a suitable substrate becomes available (17-19). Under these conditions, it is necessary for cells to organize their subcellular structures in a manner which produces the most efficient growth in order to remain competitive at both the cell and the colony level. This mode of growth is distinctly different from that observed in single cells artificially kept suspended in liquid medium, in which light, cellular contact, and gravity may come from any side, and the former two factors are generally inconsistent in magnitude (20). Under those conditions, any substance or force may be encountered from any side at any time. On a substrate, however, gravity is always applied to the cell from a constant direction, light is generally delivered from the opposite side, and once contact with other cells is made, it tends to continue for as long as the cell is attached to its substrate. Understanding the mechanisms by which cells orient and organize themselves is critical to understanding regulation of growth and metabolism.

Unlike most bacteria, cyanobacteria contain a variety of fluorescent pigments which can be observed as they are organized within the cell *in vivo*. Herein we demonstrate via confocal microscopy and the use of mutant strains of *Synechococcus* sp. PCC 7002, hereafter PCC 7002, that in addition to the well-known ability of motile cyanobacteria to display phototactic behavior (21), a phototropic mechanism exists in nonmotile species. Furthermore, we observe gravity- and substrate-specific orientation and organization of cells.

## Materials and Methods

### Growth conditions

Where substrate was used, PCC 7002 cultures were grown under 85 μmol photons m^-2^ s^-1^ cool white light from an LED panel (Huion) at 37°C on 1% agar A+ medium plates (22). *Synechocystis* sp. PCC 6803, hereafter PCC 6803, cultures were grown under 85 μmol photons m^-2^ s^-1^ cool white light from the same LED panel at 30°C on 1% agar A+ medium plates. Light was delivered to the cell either in the direction of, the direction opposite, or perpendicular to the direction of gravity. PCC 6803 wild-type strain was used; PCC 7002 wild-type (WT) and polyphosphate kinase knockout (Δ*ppk*; previously generated in the Cameron lab) strains were grown and imaged as described below. For noted cultures, a home-built spinning plate device was used to generate the effective lateral (i.e. perpendicular to natural gravity) forces ranging from 0.5 to 5 times gravity on the cells. Where PCC 7002 WT cells were grown in liquid medium, growth occurred in volumes of 100 mL in 250 mL covered Erlenmeyer flasks on a shaker set to 80 rpm, under 85 μmol photons m^-2^ s^-1^ cool white light from Philips Alto II 25W bulbs delivered from above at 37°C.

### Confocal Imaging

All confocal fluorescence images were obtained on a customized Olympus FV-3000 confocal microscopy system with Olympus UPlanXApo 60x oil immersion objective (1.42 numerical aperture) built onto an Olympus IX-83 inverted fluorescence microscope with Z-Drift Compensator and onboard excitation and imaging illumination from lasers at 405, 488, 561, and 640 nm (50, 50, 50, and 40 mW Coherent OBIS laser modules, respectively). Cells grown in liquid medium were streaked onto agar pads and inverted onto 4-well ibidi μ-Slides immediately before imaging. Cells grown on agar plates were excised with their agar in situ and carefully inverted onto 4-well ibidi μ-Slides immediately before imaging. “Phycobilin” measurements (magenta in all images) were performed using an excitation laser power of 0.02%, wavelength of 640 nm, and emission range of 650-720 nm. This wavelength excites both allophycocyanin (primary absorption) and chlorophyll and thus 9% of the yield here is from chlorophyll as well. “Chlorophyll *a*” measurements (green in all images) were performed using an excitation laser power of 0.05%, wavelength of 405 nm (which does not also excite phycobilins), and emission range of 650-720 nm. All imaging was performed at a capture time of 2.0 μs/pixel.

### Image Analysis

Images were viewed, color-coded by channel, oriented, and converted to 3D representations using ImageJ (Fiji; NIH-supported freeware). To quantify spatial resolution of pigment localization, sum pixel intensity over a plane through representative cells at each pixel of depth was obtained for each image using Fiji. These values were then plotted by pixel depth across the cell using OriginLab Origin 2018b. Contrast images by pigment were obtained by subtracting the values in the phycobilin channel from the chlorophyll channel (in Fiji) and displaying using a custom color lookup table with gradient from green (chlorophyll) to magenta (phycobilin).

### Transmission Electron Microscopy

PCC 7002 WT cells grown on agar plates were removed using a sterile nichrome loop, suspended in 100 mM mannitol in A+ medium, and frozen under pressure. These cells were then substituted with 2% OsO_4_ and 0.1% uranyl acetate in acetone and embedded in Epon/Araldite (23). Thick sections were collected on formvar-coated copper slot grids and dual-axis tilt series were obtained using an FEI Technai F30 IVEM (FEI). The IMOD software package was used to generate tomograms for figures (24,25).

## Results

Confocal imaging of cells grown in specific combinations of light, gravity, and substrate orientation were shown to produce characteristic distributions of bulk phycobilin and chlorophyll. Adhesion to substrate was found to be the determining factor in production of a “midline” region of the cell, which contains the thylakoid organizing centers and overlapping pigments under native conditions (26,27). The midline is specifically regulated by adhesion of surfaces, and cells adhered at an angle develop a tilted midline. The influence of the midline is also shown to diminish if an external force (such as gravity) is applied to cells in a direction that is not toward or away from the substrate, as seen in Figure 1B.

**Figure 1.**
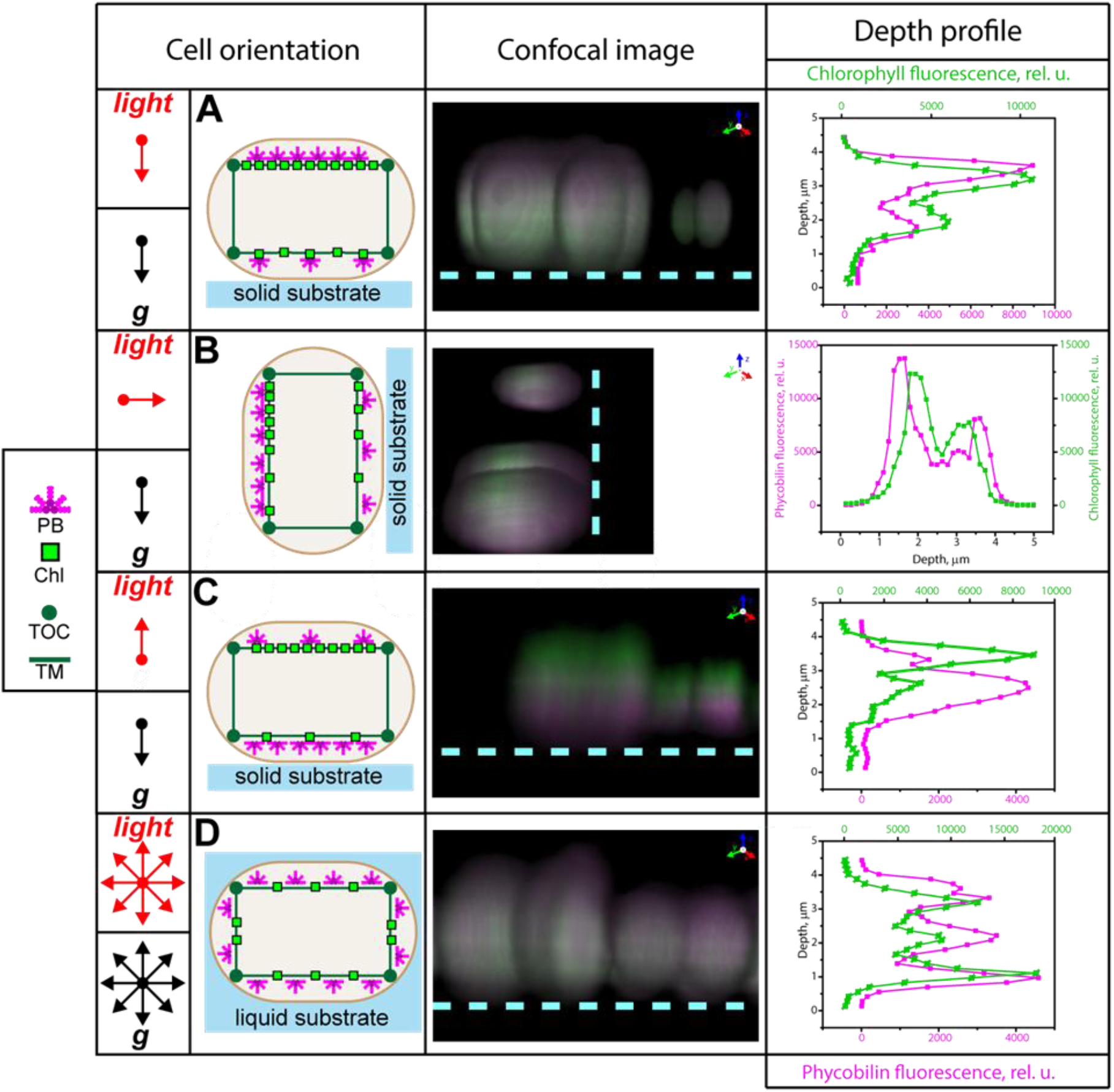
Diagrams, confocal fluorescence images (side-view) and fluorescence depth plots of PCC 7002 cells grown under (A) light from above with a standard horizontal pad orientation, (B) light from the side and a vertical pad orientation, (C) light from below with a horizontal pad orientation, and (D) light from above with liquid medium resulting in no distinct directional orientation of stimuli until cells were plated for imaging. Phycobilin (PB) shown in magenta, chlorophyll (Chl) shown in green; in diagrams, green circles are thylakoid organizing centers (TOC) and green lines are thylakoid membranes (TM). In all depth profiles, the direction of gravity is down.

While pigment is generally expressed in two bands above and below the midline of the cell, how much of each pigment is expressed in each band is determined by direction of the growth light source and the external force (in nature, gravity). Under native growth conditions (Figure 1A), both phycobilin and chlorophyll are expressed toward the upper surface of the cell, the direction opposite gravity and toward light. However, chlorophyll is overall expressed closer to the interior of the cell than phycobilin is, as well as being more evenly distributed between the two faces, and thus appears lower within images. When cells are grown on a substrate mounted perpendicular to its normal orientation, where gravity is applied along the plane of the substrate rather than into the plane, and light is applied laterally, phycobilin continues to be expressed primarily in the direction of the light, whereas chlorophyll is found on the side of the cell which faces opposite gravity. Both are still more strongly expressed opposite the substrate than toward it, though lighting the cell through near-transparent agar substrate causes reorientation of phycobilin primarily toward the substrate-bound surface of the cell. This is shown in Figure 1C, where under bottom-lit conditions an extreme division of pigment is observed; chlorophyll is expressed in the upper surface of the cell, while phycobilin is expressed in the lower. Finally, when cells are grown in liquid medium and rapidly plated and imaged, a distribution of domains similar to that observed by Vermaas et al. (28) in PCC 6803 is observed, wherein pigment is expressed in four surface domains (shown as three in the depth plot; the laterally oriented domains overlap and the midline feature is lost), and there is little overall separation of phycobilin and chlorophyll domains. The PCC 7002 morphology observed in Figure 1C can also be induced in PCC 6803, as shown in Figure S1, by growth under the same conditions.

While gravity is inevitable under terrestrial conditions, other physical forces may be applied on the cell. To examine the effects of artificial gravity on cellular control of chlorophyll expression, a spinning plate device was constructed to apply forces in excess of natural gravity (Figure 2A). Light was delivered uniformly from above the spinning plate while cells were grown on it. At the single-cell level, phenotypes similar to that demonstrated in Figure 1B, but with less extreme distribution of chlorophyll, were observed. However, differences in pigment distribution between nearby cells, especially cells forming a single microcolony, were also observed, producing apparent local gradients.

**Figure 2.**
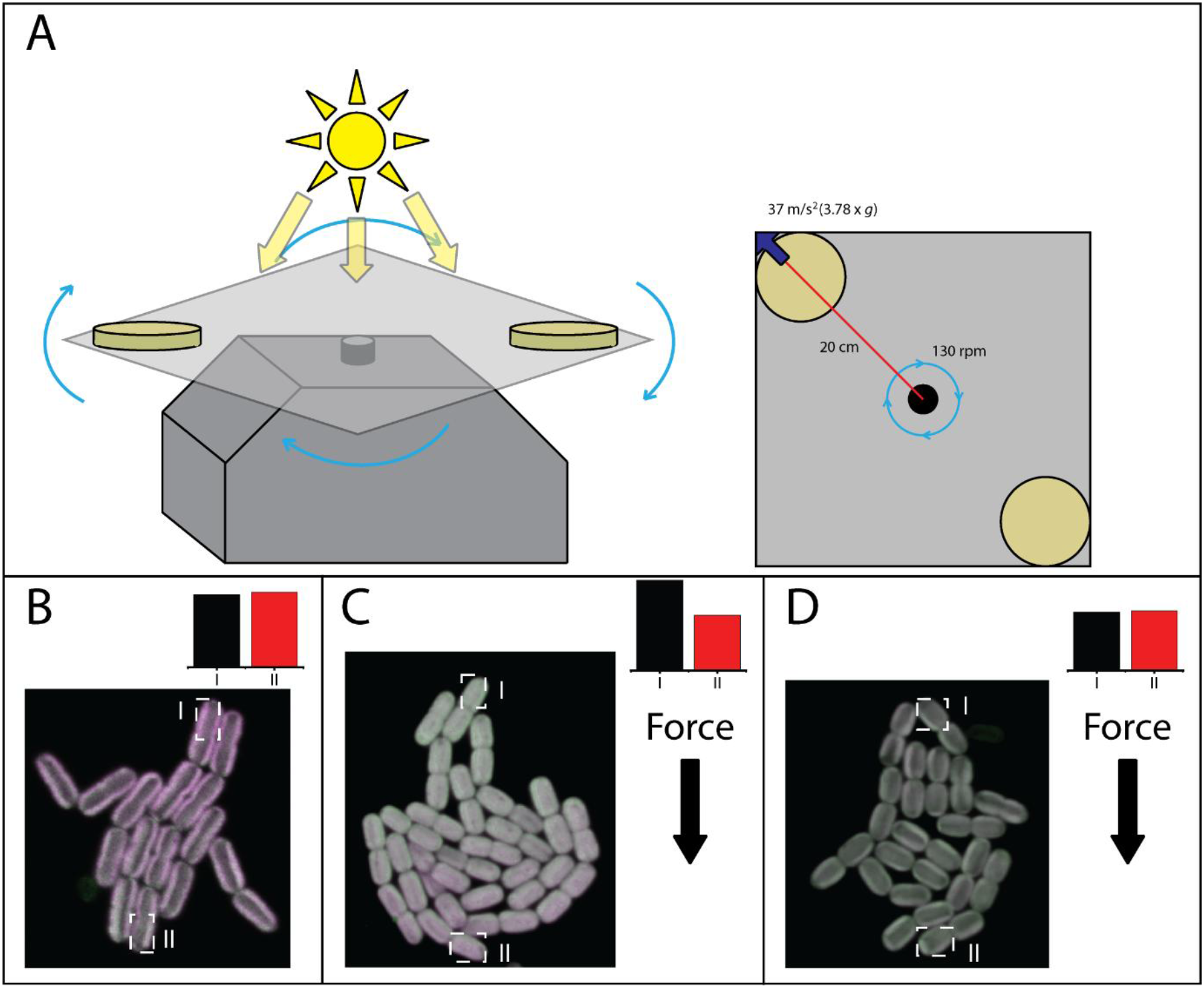
Spinning plate growth apparatus and effects of lateral force on the cell growth. (A) Growth stage allowing application of lateral force to cells showing positions of growth plates and standard force applied. (B) Example WT microcolony entering the 32-cell stage grown without applied lateral force. (C) WT microcolony exiting 32-cell stage following growth under a lateral force of 3.78x*g*, where the applied force pointed down the page. Pigment gradient characteristic of lateral force is observed. (D) Microcolony of the Δ*ppk* mutant at the 32-cell stage grown with a lateral force of 3.78x*g*, where the applied force pointed down the page. No pigment gradient is observed. The insets in (B), (C), and (D) show the chlorophyll:phycobilin ratios (in relative units) at the ends of the microcolony relative to the force vector (except in (B), where there is no lateral force applied).

A top-down view of a conventional PCC 7002 microcolony (29), grown under the same standard conditions as described for Figure 1A, is shown in Figure 2B. Cells were found reliably forming this “hand”-shaped morphology in microcolonies at the 16- and 32-cell stages of growth in order to minimize mechanical stress due to confinement. Under a strong external force, however, a “comet” morphology (Figure 2C) is adopted during these stages as a result of cells being pulled by the external force. The extent to which this phenomenon is observed is dependent on the orientation of the progenitor cell of the microcolony, which will determine the orientations of its daughter cells and therefore the extent to which cell binding to the substrate is affected by the laterally applied force and how this interacts with the normal mechanical pressure of adjacent cells (29). In wild-type cells, this morphology allows easy observation of a pigment gradient across the entire microcolony, wherein cells opposite the direction of external force express considerably more chlorophyll and less phycobilin, while cells closer to the direction of the force produce more phycobilin at the expense of chlorophyll. However, in the Δ*ppk* mutant strain (Figure 2D), which lacks polyphosphate (30-32), no such gradient is observed under applied external force. Aside from decreased overall phycobilin due to the lack of polyphosphate, the individual cells are phenotypically similar to those in Figure 2B, though the overall colony morphology is still visibly altered. Polyphosphate is therefore necessary for force-induced pigment redistribution.

While cells are visibly capable of responding to interaction with other cells under lateral force, they are also capable of responding to other cells under natural gravity conditions, provided the interaction is in the direction of gravity. Figure 3A and 3C show top and side views of a slice of a multi-layer cell colony in which a pigment gradient can be observed across all vertical layers of cells. While all cells contain both chlorophyll and phycobilin, distributed normally within the cell, the ratio of the two pigments steadily shifts from strongly favoring chlorophyll on the bottom (substrate) surface to expressing mostly phycobilin near the top (facing the light). Total pigment expression is also higher in cells closer to the bottom of the colony, which is consistent with shading from higher up; cells in higher levels have proportionally less overall pigment and the uppermost levels distinctly contain less pigment than monolayer cells. However, application of lateral force causes a significant reorganization of the pigment distribution, as seen in Figure 3B and 3D. In addition to some degree of deformation of the colony itself, chlorophyll expression is shifted toward one end of the colony. The applied force is stronger than gravity, which results in cells at the interior producing proportionally elevated phycobilin more consistent with upper-level cells in a normal colony. Figure parts 3E and 3G reveal the domains present in these two types of colonies.

**Figure 3.**
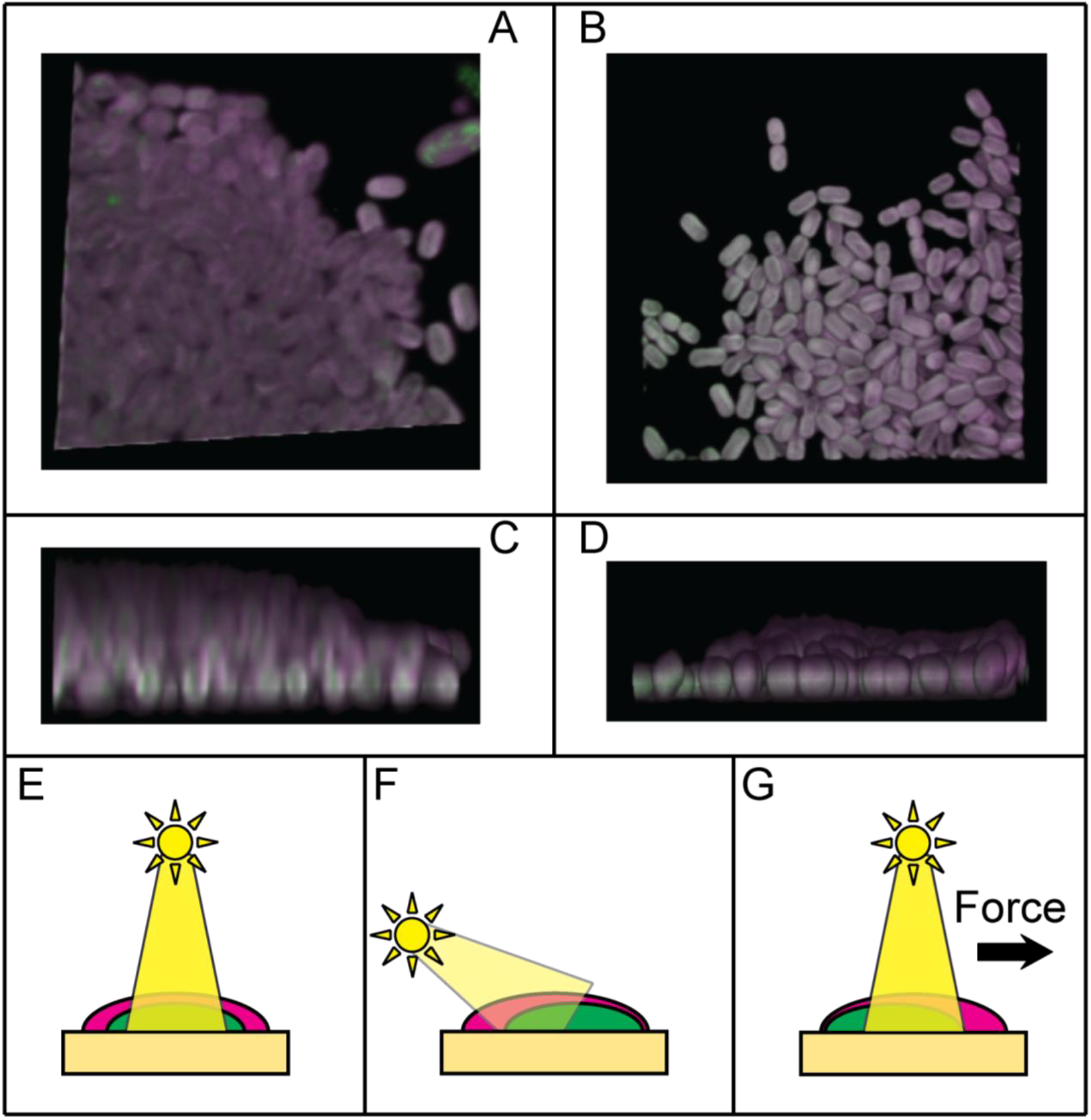
Top row: Top views of multi-layer WT colonies grown under (A) normal conditions without applied external force and (B) grown under a lateral force of 3.78x*g*. Middle row: (C) and (D) are side views of the same colonies in (A) and (B), respectively. Bottom row: Diagram of expected colony-scale pigment distribution under various growth conditions: (E) overhead sunlight (as in (A) and (C)); (F) angled “morning”- or “evening”-type sunlight directionality; (G) applied lateral force to exceed gravity (as in (B) and (D)).

Cells are also capable of responding to the presence, and pigment distribution state, of nearby cells with which they do not have physical contact. The microcolonies visible in Figure 4A show a considerable proportional overexpression of chlorophyll when farther from gravity and a similar increase in phycobilin lower on the growth plate, despite that these microcolonies are separated by several microns. Gradients exist both within individual microcolonies and across the overall group. This implies that the cells are able to detect the presence of other cells via a signaling molecule which can diffuse through the agar substrate. However, as seen in Figure 4B, when these cells are starved of phosphate, they are unable to detect the presence of other cells above them and no significant pigment gradient can be observed.

**Figure 4.**
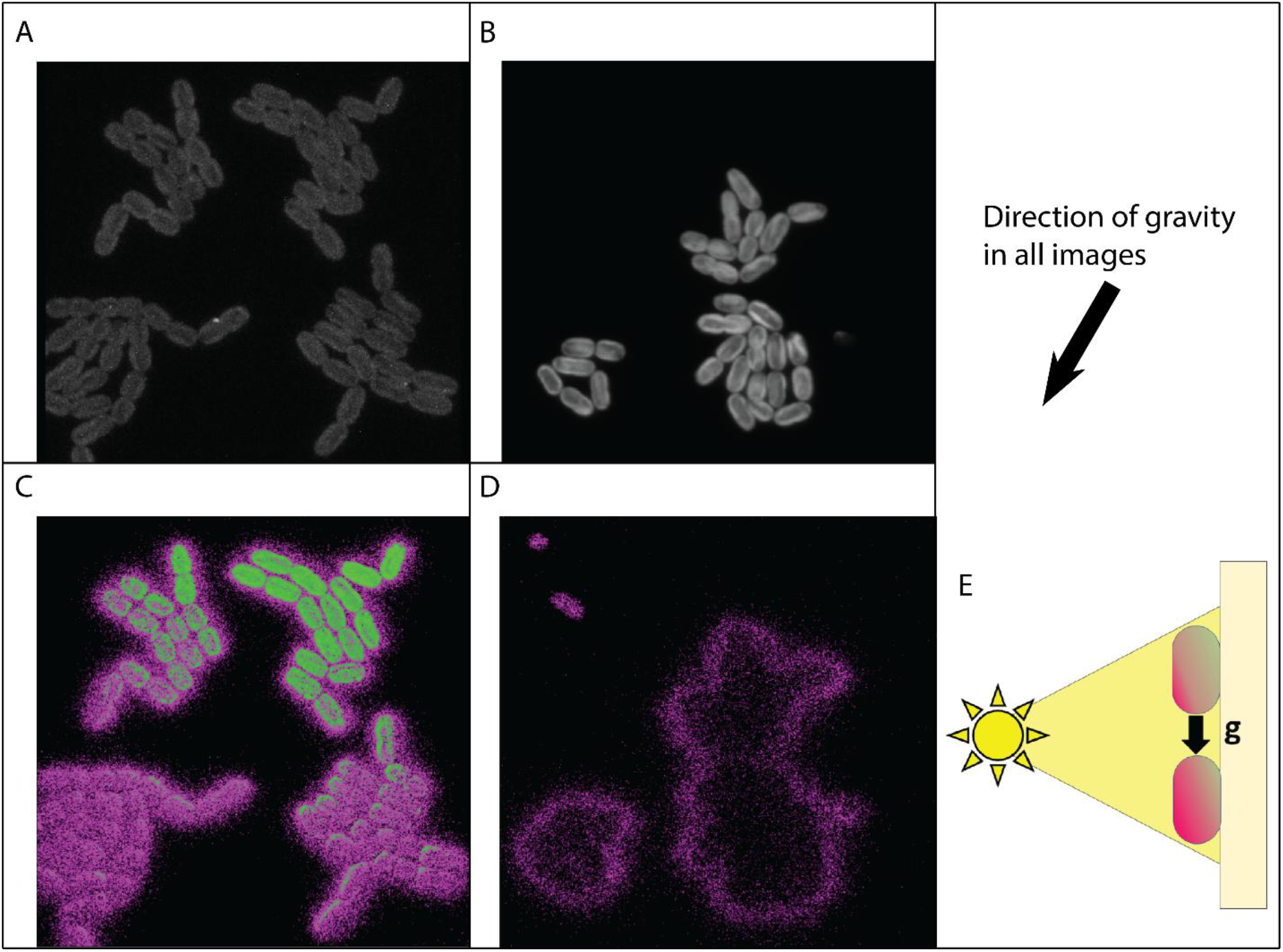
(A) A confocal microscope image (shown from the top view) of microcolonies of PCC 7002 demonstrating pigment gradients both within and between colonies’ distinct pigment regions. This imaging was done within a multi-layer cell colony after growth immobilized on 1% A+ agar with side lighting and the plate oriented toward the light (direction of gravity is marked). (B) Microcolonies under otherwise identical conditions grown without phosphate until near-starvation state. (C) Difference between chlorophyll (green) and bulk phycobilin (magenta) channels of the image in (A). (D) Only scattering visible in a similar gradient image of (B) due to negligible difference in pigment expression across the colony. (E) Schematic of intracellular pigment distributions as determined by position on a vertically oriented pad with light delivery toward the face of the pad, perpendicular to gravity.

To elucidate the mechanism by which polyphosphate might be involved in responding to gravity, transmission electron microscopy was employed. PCC 7002 produces multiple types of polyphosphate bodies, including the classical spherical structures, which are very large relative to the size of the cell in PCC 7002 (Figure S2), and a stalked structure consisting of a small tether at the thylakoid organizing center and a larger domain extending into the cytoplasm. The stalked structure and the large polyphosphate body may fuse together (Figure 5) to form structures large enough to be observed via visible-light transmission-mode microscopy. While the weight of the large polyphosphate bodies in this cyanobacterium may be adequate on its own to direct a cell response by pressing on the thylakoids as in Figure S2, if such a structure were attached to the thylakoid organizing center in a manner as in Figure 5, such a structure would apply an outsize gravitational torque at the organizing center, which could itself apply a mechanical action on the thylakoids, prompting a response needed to balance the polyphosphate body.

**Figure 5.**
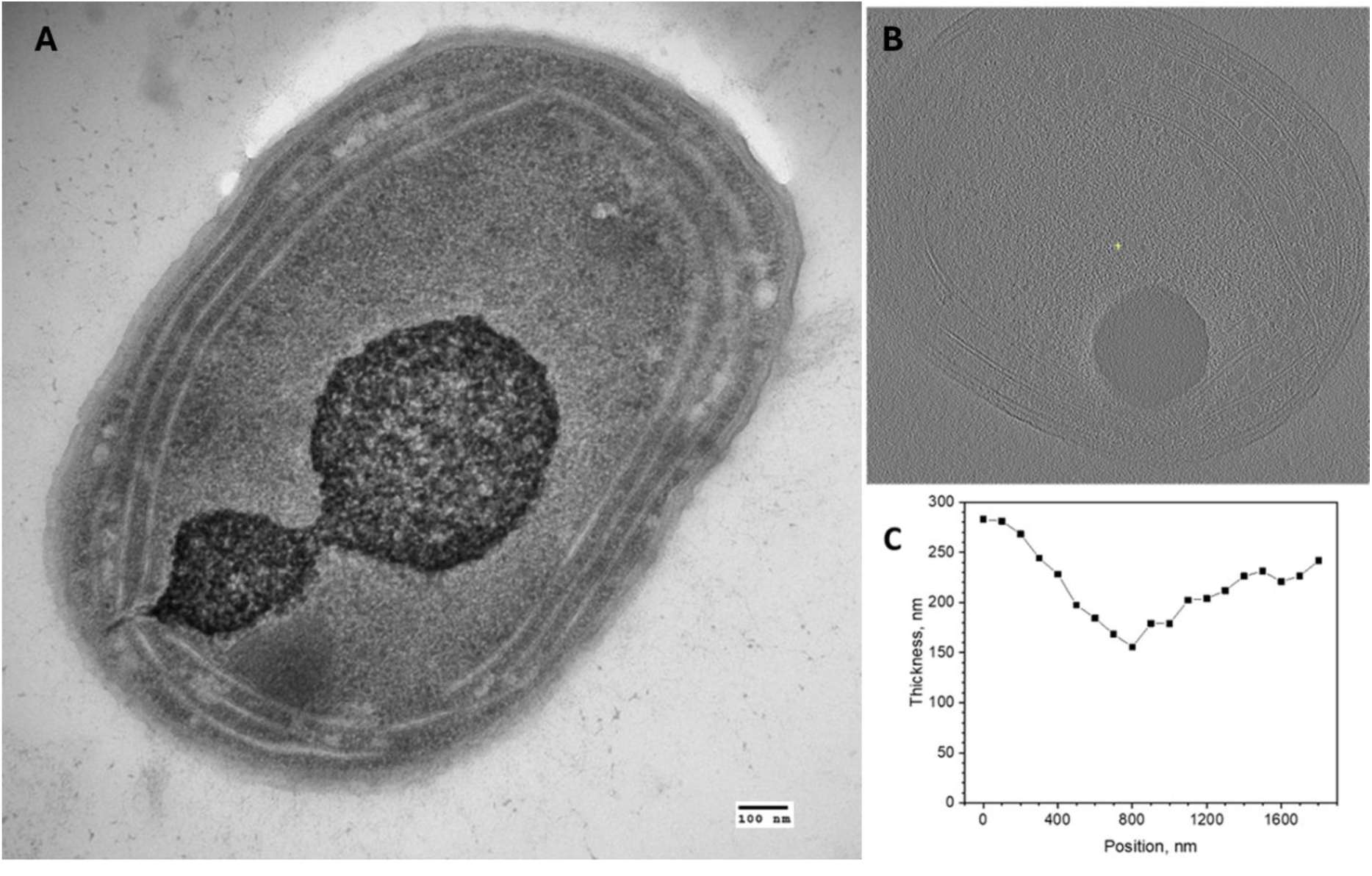
(A) Cryo-electron micrograph of a section through a PCC 7002 cell showing a thylakoid organizing center (small gap in thylakoid membranes) at which is attached a double polyphosphate body. (B) Cryo-electron microscopy image of PCC 7002 as generated using the IMOD visualization software, showing displacement of thylakoid membrane by a polyphosphate body (bottom center). (C) Plot of the distance in nanometers between the center of the innermost leaf of thylakoid membrane and the cell wall at 100 nm intervals in (B), moving clockwise away from the thylakoid organizing center (gap in membranes, at right) and passing the polyphosphate body along the bottom of the cell. The total thickness of the thylakoid region is compressed 45% at the center of the polyphosphate body.

## Discussion

### Physical basis of gravitropic response

It is instructive to first note that even within the colloidal science perspective, when treating entire cyanobacteria and polyphosphate bodies within them as inanimate colloidal objects (dead cells), gravity plays an important role in controlling behavior like sedimentation to substrates and the tendency of orientation of both individual cells and trichomes formed by their assembly relative to the underlying support substrate assumed to be perpendicular to the gravity force direction (33,34). The gravity forces and torques acting on most cyanobacteria with individual cell dimensions in the micrometer range easily overcome the strength of thermal fluctuations, causing sedimentation and parallel-to-substrate alignment (33). This primitive perspective of anisotropic particles is influenced by gravity at different levels of assembly, from trichomes to biofilms, but is insufficient to explain effects discussed in the introduction, where our present study provides some insights. For example, it is the presence of polyphosphate bodies, not bulk phosphate, which allows the cell to modify its pigment distribution in response to a force being applied to it. The Δ*ppk* mutant cells in Figure 2D were grown on sufficient phosphate so that growth rate is not noticeably affected, but are unable to form a pigment gradient consistent with the force being applied. Accordingly, it is the presence of the polyphosphate body, not the available phosphate level, which enhances the gravitropic response.

The mechanism by which the polyphosphate body is used to determine direction of gravity may rely on the movement of the polyphosphate body, or pressure on its surrounding protein matrix, within the cell itself; or it may be due to the force exerted by the overall cell (moving with the polyphosphate body) on its exopolysaccharide matrix. Loss of polyphosphate results in significant loss of the exopolysaccharide matrix (35), although cells still appear to be able to determine the direction of substrate and form midlines accordingly. This response is rapid enough to favor an internal mechanism not reliant on the building of complex structures such as an exopolysaccharide matrix prior to pigment redistribution. The likely reason that the substrate is playing a role in determining the pigment distribution is that almost any substrate under natural habitat conditions would be opaque and, thus, light would not generally arrive from this direction. This could have been utilized during its evolution to allow the cell to establish a long-term pigment distribution to optimize use of incoming light.

The use of an internal object with density considerably higher than the surrounding cytoplasm and size sufficiently smaller than the cell itself to allow some degree of movement is found in analogous systems across the tree of life. In animal cells, the cytoskeleton and its interactions with dense organelles are involved in sensing the direction and force of gravity being applied (9,36). Some fungal lineages use sedimentation of protein crystals (octin) as a gravity-sensing mechanism (10), as one of multiple similar mechanisms extant in fungi (37). In plant cells, perhaps the best-understood gravity-sensing system developed in nature, amyloplast sedimentation within specific cells (statocytes) is used to direct both root and shoot growth (8,38). The amyloplast is similar in size as a fraction of the total volume of the cell to a standard polyphosphate body in PCC 7002 (39,40). While the amyloplast is roughly 1.5 times the density of the cytoplasm through which it moves, the plant cell is much larger and less susceptible to Brownian motion than a bacterial cell; however, polyphosphate bodies are considerably denser than amyloplasts, 1.8-2.1 times the density of overall bacterial cytoplasm (41). Treating the bacteria as colloidal particles within an aqueous environment, it is possible to calculate whether the bacteria themselves within medium or the polyphosphate bodies within the cytoplasm are susceptible to sedimentation using the following equation for gravitational length (42):

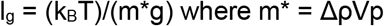

The bacterial cells used here averaged 4.5 cubic microns in volume, while the polyphosphate bodies of interest had volumes between 0.0082 and 0.11 cubic microns. A typical gram-negative cell density is 1.11 g/ml (43), while its cytoplasm has an aqueous solvent (1.0 g/ml) and polyphosphate bodies in cyanobacteria are primarily magnesium polyphosphate, 2.1 g/ml, but may be comprised of heavier calcium and magnesium polyphosphates as well, reaching up to 3.5 g/ml. Sedimentation of the cell itself thus does take place, but not quickly or universally, as treating a suspension of cells as a colloid finds its gravitational length to be 0.88 microns, which is smaller than the shorter dimension of PCC 7002 cells (and thus its colloidal diameter) by a relatively small margin-this dimension is 1.5-2.0 microns in almost all cells. A colloidal particle must have a gravitational length smaller than its diameter to precipitate, i.e. a precipitation ratio of less than 1 (44). As the cell’s ratio is quite close to this value, it would conceivably be able to avoid sedimentation by degrading dense internal polymers such as polyphosphate to reach a density of 1.065 g/ml or producing extracellular polymeric substances (EPS) with lower density to increase the particle diameter or lower the net density of the particle. Indeed, both of these phenomena are observed in other cyanobacteria (45,46).

**Table 1.** Gravitational length and object diameter for PCC 7002 and its polyphosphate bodies.

|  | PCC 7002 | Polyphosphate body |
| --- | --- | --- |
| Gravitational length $l_g$, $\mu\text{m}$ | 0.88 | 3.6-48 |
| Effective diameter, $\mu\text{m}$ | 1.5-2.0 | 0.25-0.6 |
| Precipitation ratio | 0.44-0.58 | 6.0-190 |
| Gravitational torque | $6.50 \times 10^{-20}$ J | $2.45 \times 10^{-21}$ J |

While the bacterial cell can be expected to sediment on its own under most conditions, even its largest spherical polyphosphate body is too small to overcome Brownian motion, though the extended structure, almost a micron long, is nearly large enough to precipitate. A polyphosphate-based mechanism may rely on an added torque effect on the “stalk” to the thylakoid organizing center, as the polyphosphate body is approximately three orders of magnitude more massive than the small polyphosphate intrusion at the thylakoid organizing center. This results in a gravitational torque of 2.45*10^-21^ J (N*m), assuming a polyphosphate body of the size shown in Figure 5A, which is weaker than Brownian motion (thermal torque) at the growth temperature (4.18*10^-21^ J)(47-49). The cell itself is large enough to have a gravitational torque about its center exceeding Brownian motion by over an order of magnitude. Thus, it is not clear that the pull of gravity on the polyphosphate body is a mechanism that can reliably be utilized by the cell, and indeed large non-stalked polyphosphate bodies are sometimes observed freely floating within the cell, though typically these are smaller, below 400 nm in diameter. However, it is also possible that the polyphosphate body itself simply alters local chemistry in a manner that favors metabolic and translational regulation in conjunction with the cell’s own sedimentation process or internal torque by providing a fixed point.

### Broader implications of polyphosphate in gravitropic behavior

Polyphosphate is known to be necessary for EPS production and biofilm formation in several bacterial lineages, including other cyanobacteria (50-52). There is also some precedent for bacteria using polyphosphate in what is unequivocally a gravitactic response; sulfur-oxidizing bacteria in the Black Sea have evolved a mechanism whereby they build up massive polyphosphate granules (large enough to escape Brownian motion, several times more massive than those observed here) in order to sink to sulfur-rich regions of the water column and dismantle these granules to become positively buoyant and rise to oxygen-rich layers (53). Additionally, magnetotactic bacteria, which can orient horizontally by the Earth’s magnetic field, accumulate large polyphosphate bodies, likely to add a vertical dimension to their ability to orient themselves and increase stability (54). As gravity response mechanisms have evolved independently several times (55), these may not be directly related to that found in the cyanobacterial lineage. However, cell positioning and response to gravity using this mechanism may be more fully realized in filamentous cyanobacteria, which due to their considerably larger size (filaments vs. cells) can much more easily overcome Brownian motion assuming a similar density. These cyanobacteria can also overcome gravitational sedimentation in solution by producing EPS (56) to increase their flat surface area and lower the average density of the overall particle to below that of water, as well as by forming mats and rafts with other filaments (57) and generating gas vesicles to become buoyant (58).

### Interaction of the phototropic and gravitropic responses

The likely cause of the existence of two separate regulatory mechanisms for chlorophyll and phycobilin expression within the cell is the need to respond to changing light conditions during the diurnal cycle. For much of the day, the sun is not directly overhead, and at these times, a cyanobacterial colony will experience relatively low light levels on average. Less light reaches the Earth’s surface at morning and evening; objects are more likely to cast shadows onto the cells at these times; and finally, the light has to pass through more cells to reach most points in a mature colony or biofilm than it would if delivered from directly overhead. Under these conditions, the limiting factor in cell growth is usually the supply of reductant (59,60), which means that for maximum growth rate, as much light as possible should be directed to photosystem II (PSII) rather than photosystem I (PSI). Cyanobacteria already have a very low PSII:PSI ratio as compared to other photosynthetic clades (61), which is mitigated by expression of large, effective phycobilin antenna complexes. Orienting PSII and the associated antenna complexes to capture as much light as possible permits maximal growth under light-limited conditions.

Conversely, when the sun is directly overhead, an excess of light is available to much of the cyanobacterial colony. Under this condition, phycobilins are dissociated from PSII and serve as a photoprotectant, dissipating incoming light as heat and avoiding destructive recombination (62). As for PSI, with which most of the chlorophyll in the cell is associated (63), it is advantageous to have this photosystem oriented opposite the direction of gravity when the sun is directly overhead because it can then perform cyclic electron flow (PSI-CEF) as an added photoprotective mechanism (64). Under very low light intensities (i.e. when the sun is not directly overhead, or deep in the colony), it is also beneficial to express high levels of PSI because PSI-CEF captures some energy with less risk of recombination producing dangerous radicals than is the case in PSII under low light conditions. For related reasons, PSI is longer-lived than the frequently replaced PSII (65). This two-regulator system, thus, effectively takes advantage of both the light-harvesting and the photoprotective qualities of the two major pigments at the cell and the colony scale.

The importance of gravity in cyanobacterial colony development and photosynthetic regulation has been noted in a recent study which found impaired growth under microgravity (66), which was attributed to the formation of an uncharacteristically thick boundary layer which trapped oxygen and inhibited photosynthesis. In addition to the misalignment of photosynthetic pigments which we would expect under these conditions based on the results herein, it stands to reason that a poorly directed external polysaccharide matrix, which is known to be reliant on the presence of polyphosphate under normal conditions (35), would occur if extant polyphosphate cannot be used to detect the direction of gravity.

## Conclusion

We have demonstrated that gravitropism interplays with phototropism to significantly define cyanobacterial behavior at levels that go well beyond the simple influences of gravity-driven sedimentation to substrates or gravitational torque-influenced alignment with respect to the substrates. Intracellular distribution of chlorophyll and phycobilin in PCC 7002 is regulated by three factors: 1) the side of the cell which is bound to the substrate; 2) the orientation of the cell relative to gravity force direction; and 3) the direction from which the cell is lit. The substrate-proximity-defined direction controls where in the cell thylakoid domains are oriented, gravity is used to determine where to express chlorophyll, and light direction controls phycobilin expression. Cells are capable of forming multicellular pigment gradients to optimize growth, though pigment gradient formation is dependent on the presence of polyphosphate bodies. PCC 7002 naturally sediments over time and this sedimentation is necessary for the development of spatially resolved internal physiological domains. Our findings could have wide-ranging implications in diverse research fields, spanning from the designs of new forms of topological active matter (67) to the designs of habitats for the colonization of extraterrestrial bodies (68).

Our study highlights the need for additional systematic research to further elucidate the role of gravity and illumination directionality in terms of various aspects of cyanobacterial behavior, including individual cells, trichomes and larger cell communities. Specifically, the effects of a complete lack of directional force and the role of a light source of inconsistent direction or intensity remain to be determined.

## Acknowledgments

This study was financially supported in part by the U.S. Department of Energy (DOE) DE-SC0019306 (to J.C.C.). We thank Nicholas Hill for originally generating the Δ*ppk* strain, Janet Meehl for assistance generating electron micrographs, and Keita Richardson and Kristin Moore for constructive feedback.

