## Supplemental Figure 1 for "Interactions of Phototropism and Gravitropism in Cyanobacteria"

**Supporting Information**

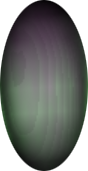

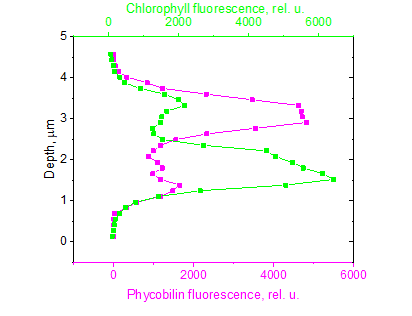

**Figure S1**. (A) Confocal microscope image (shown from side view) of a cell of *Synechocystis* sp. PCC 6803 demonstrating distinct pigment regions after growth immobilized on 1% A+ agar with bottom lighting. (B) Depth profile of the cell as in Figure 1.
